# Mutual interaction between Visual Homeostatic Plasticity and Sleep in Adult Humans

**DOI:** 10.1101/2021.03.31.437408

**Authors:** Danilo Menicucci, Claudia Lunghi, Andrea Zaccaro, Maria Concetta Morrone, Angelo Gemignani

## Abstract

Sleep and plasticity are highly interrelated, as sleep slow oscillations and sleep spindles are associated with consolidation of Hebbian-based processes. However, in adult humans, visual cortical plasticity is mainly sustained by homeostatic mechanisms, for which the role of sleep is still largely unknown. Here we demonstrate that non-REM sleep stabilizes homeostatic plasticity of ocular dominance induced in adult humans by short-term monocular deprivation: the counter intuitive and otherwise transient boost of the deprived eye was preserved at the morning awakening (>6 hours after deprivation). Subjects exhibiting a stronger boost of the deprived eye after sleep had increased sleep spindle density in frontopolar electrodes, suggesting the involvement of distributed processes. Crucially, the individual susceptibility to visual homeostatic plasticity soon after deprivation correlated with the changes in sleep slow oscillations and spindle power in occipital sites, consistent with a modulation in early occipital visual cortex.

## Introduction

Neural plasticity is an intrinsic property of the nervous system underlying the ability to change in response to environmental pressure. Learning and memory processes plastically and continuously encode new neuronal information during wakefulness, but consolidation mechanisms often extend into sleep^1^. During non-REM (NREM) sleep, replay and pruning processes as well as connectivity rearrangements associated with specific neuronal activity patterns involving GABAergic modulation^2^ play a fundamental role in plasticity consolidation.

Two hallmark rhythms characterize the background of on-going oscillations of NREM sleep: slow wave activity (SWA, 0.5-4 Hz) and sigma band (σ, 9-15 Hz). A fundamental contribution to the lower frequency bound of SWA is due to Sleep Slow Oscillations (SSOs), an EEG pattern that corresponds to the alternation between periods of neuronal membrane depolarization and sustained firing (up states) and periods of membrane hyperpolarization and electrical silence (down states) whereas power in the sigma rhythm is provided by the sleep spindles: waxing and waning wave packages that spread throughout the thalamocortical system^3^. Sleep slow oscillations and sleep spindles have been consistently associated with synaptic plasticity, replay and memory consolidation^1, 4, 5^. SSOs allow spike-timing-dependent-plasticity, while replay occurs via cortico-thalamo-cortical interactions that are made effective through thalamus-cortical synchronization in the sigma band: the sleep spindle is the full-fledged expression of this mechanism^6^. Several studies have investigated the role of slow and spindle oscillations in episodic and procedural memory^7, 8^, while the contribution of these processes in sensory plasticity has yet to be assessed.

Growing evidence indicates that the adult human visual system might retain a higher degree of plasticity than previously thought^9, 10^. Some degree of Hebbian plasticity is retained in adulthood and mediates visual perceptual learning^11^ as well as visuo-motor learning^12, 13^. The residual Hebbian plasticity in adult involves changes in neural processing occurring at multiple levels of visual (perceptual learning^14^) and visuo-motor^15^ processing, particularly at associative cortical level, and are consolidated both by NREM^12, 13^ and REM^16^ sleep. While sleep-dependent Hebbian plasticity has been shown in thalamo-cortical circuits in rodents visual system following prolonged exposure to a novel visual stimulus ^17^, Hebbian plasticity of ocular-dominance in the primary visual cortex (V1) is very weak or absent in primates after closure of visual critical period.

Ocular dominance plasticity^18–20^ is an established model of sensory plasticity in V1 in vivo, which is observed after a period of monocular deprivation (MD) ^20, 21^. Ocular dominance plasticity is maximal during development, when it is mediated both by Hebbian and by homeostatic plasticity, which have different functional outcomes. Hebbian plasticity stabilizes the most successful inputs in driving neural activity, and consequently the deprived eye loses the ability to drive cortical neurons^22^. On the other hand, homeostatic plasticity upregulates the neuronal response gain of the weakened deprived eye. During development, Hebbian and homeostatic mechanisms work hand in hand and their relative strength depends on timing and monocular deprivation duration ^23, 24^. The homeostatic ocular dominance plasticity is preserved through the life-span: in adult humans, recent studies have shown that short-term MD (2-2.5h) unexpectedly shifts ocular dominance in favour of the deprived eye ^25–27^. This counterintuitive result, consistent with homeostatic plasticity, is interpreted as a compensatory adjustment of contrast gain in response to deprivation. The deprived eye boost is observed also at the neural level, as revealed by EEG^28^, MEG^29^ and fMRI^30, 31^ and, importantly, it is mediated by a decrease of GABAergic inhibition in the primary visual cortex^32^. The effect of short-term MD decays within 90 minutes from eye-patch removal^25, 27^. However recent evidence from a clinical study in adult patients with anisometropic amblyopia, shows that repeated short-term deprivation of the amblyopic eye can promote the long-term recovery of both visual acuity and stereopsis^33^, suggesting that the effect of short-term MD can be consolidated over time. In adult amblyopic patients, the classic occlusion therapy ^34^, which consists in the long-term deprivation of the non-amblyopic eye and relies on Hebbian mechanisms, is much less effective than the inverse occlusion approach (234 hours of traditional occlusion per 1 line of visual acuity improvement ^35^ vs. 12 hours of inverse occlusion per 1.5 lines of visual acuity improvement ^33^). Inverse occlusion involves the short-term deprivation of the amblyopic eye, relies on the homeostatic plasticity mechanisms described above and is consolidated for up to one year^33^. Understanding the role of sleep for the maintenance of visual homeostatic plasticity induced by short-term monocular deprivation is therefore a clinically relevant and timely question.

Evidence from animal models shows that NREM sleep is necessary to consolidate Hebbian mechanisms during ocular dominance plasticity within the critical period^36–38^. However, it is still largely unknown whether sleep has similar effects after the closure of the critical period in the visual cortex. In addition, it is still unknown whether sleep can modulate homeostatic plasticity induced by MD ^25–27,29,30,39–44^

There are several mechanisms that are shared between homeostatic plasticity and sleep, indicating a possible interaction between the two phenomena. Homeostatic plasticity is based on GABA-dependent mechanisms^45, 46^ that alter the excitation/inhibition balance and appear to be analogous to the factors that modulate the expression of slow-wave sleep ^47^. More generally, plasticity induced by learning does change local cortical mechanisms that are stabilized by sleep, and the stabilization relies upon hippocampal activity during NREM in several experimental models including sensory memory^48–51^. The potential hippocampal involvement can be traced by measuring sleep spindles that invade the cortex from this structure during NREM sleep in order to support memory consolidation.

Here we investigate the effect of NREM sleep on visual homeostatic plasticity in adult humans, both analysing sleep features at the occipital and the prefrontal cortex levels.

## Results

We assessed visual cortical plasticity by measuring the effect of short-term (2h) of monocular deprivation (MD) on ocular dominance measured by binocular rivalry^52–54^ in adult volunteers. In the experimental night (Monocular Deprivation Night, MDnight), MD was performed in the late evening and was followed by 2h of sleep, during which high-density EEG was recorded (Figure 1A). At this night awakening, ocular dominance was assessed again, and then participants went back to sleep until the morning (4-5 hours of additional sleep). For the control condition (Control Night, Cnight), the same participants underwent an identical protocol, but without performing MD (Figure 1B).

**Fig 1.**
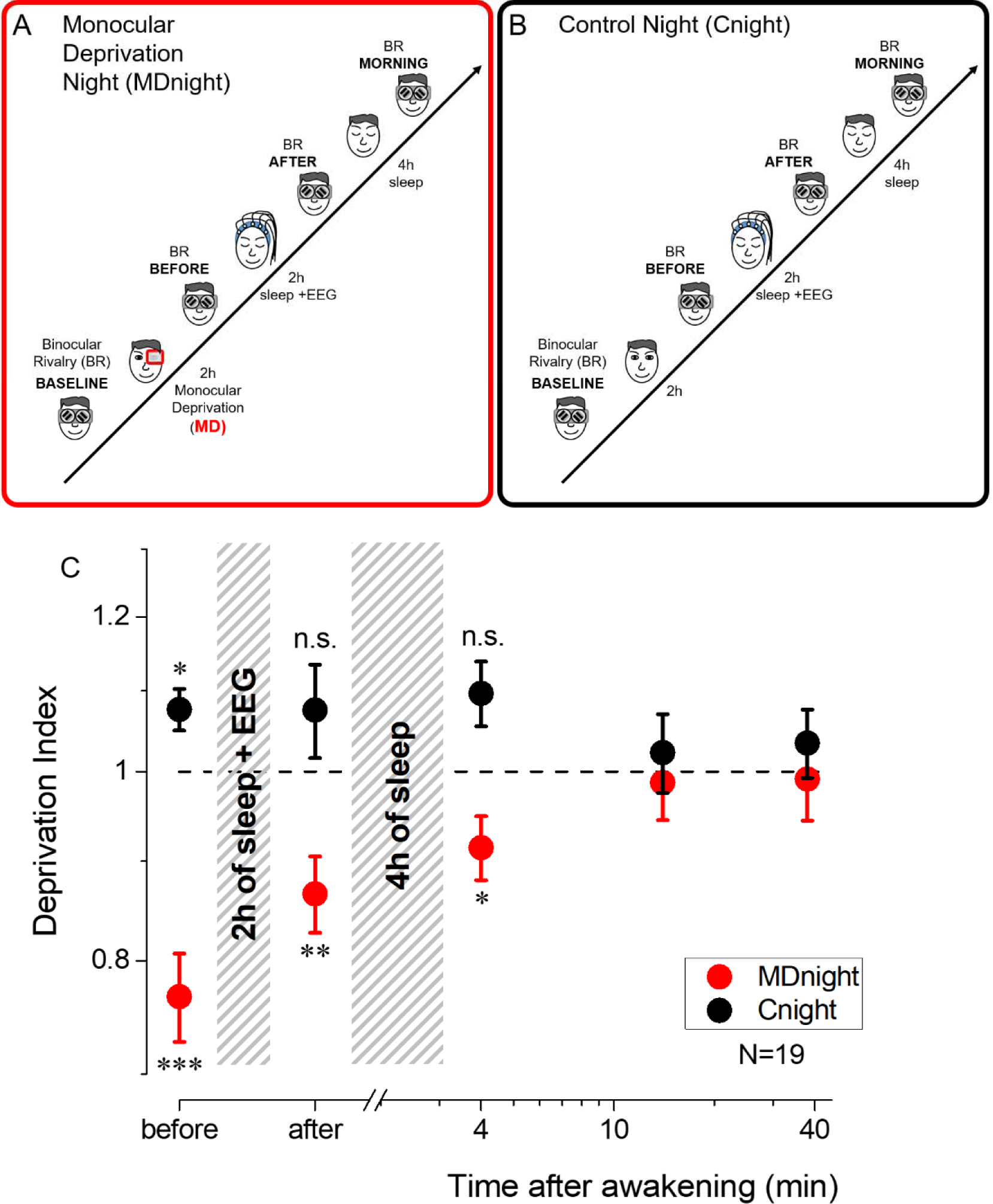
Experimental paradigm and monocular deprivation effect before and after sleep. **A**) Diagram of the experimental paradigm for the MD night (MDnight) condition. Ocular dominance was measured by means of binocular rivalry (BR) before and after 2h of monocular deprivation (MD). Afterwards, participants went to sleep while their EEG activity was recorded with a 128 electrodes system. Binocular rivalry was measured after 2 hours of sleep and in the morning, after 4 additional hours of sleep. B) same as A, but for the control night (Cnight). The experimental procedure was the same as the MDnight, except that participants did not underwent monocular deprivation **C**) The MD effect (deprivation index) measured before, after 2 hours of sleep and at morning awakening, occurring after 4 additional hours of sleep, in the MD (red symbols) and control (black symbols) nights. Error bars represent 1±SEM. Asterisks indicate the significance level (t-test of individual time-points against the value 1) after correction for multiple comparisons: ***p<0.001, **p<0.01, *p<0.05 or non-significant (n.s.).

Consistently with previous reports^25, 26^, short-term MD shifted ocular dominance in favour of the deprived eye (Figure 1C, red symbols, repeated measures ANOVA, F (4,72) = 6.7, p < 0.001, η2 = 0.27): the deprivation index (DI) was significantly altered just after eye-patch removal (mean DI ± SE = 0.77 ± 0.04, two-tailed, one sample t-test (t(18) = -5.41, p_fdr = 0.00017, Cohen’s d = 1.24).

Importantly, this form of homeostatic plasticity was maintained after two hours of sleep (mean DI ± SE= 0.87 ± 0.04, two-tailed, one sample t-test t(18)=-3.54, p_fdr=0.0046, Cohen’s d = 0.81) and during the first 8 minutes after morning awakening (mean DI ± SE = 0.91 ± 0.04, two-tailed, one sample t-test t(18)=-2.55, p_fdr=0.02, Cohen’s d = 0.59), that is about 6-7 hours after eye-patch removal. All those measurements are referred to the baseline condition taken just before the deprivation. No consistent changes in ocular dominance were observed in the control night (Figure 1C, black symbols, repeated measures ANOVA, F (4,68) = 0.97, p = 0.43, η2 = 0.05), when calculating the same index for measurement at the same time but without monocular deprivation.

Exploratory post-hoc analyses revealed however that in the control night the deprivation index measured before sleep, compared to the same measurement performed 2h before, was significantly larger than one (mean DI ± SE = 1.08 ± 0.03, paired-samples t-test t(17)=-2.7, p_fdr=0.045, Cohen’s d = 0.62), favouring the non-dominant eye. This might reflect a transient form of adaptation or implicit learning due to the repetition of the binocular rivalry (BR) test^55^, suggesting that the effect of monocular deprivation may be overall underestimated.

That the effect of deprivation is maintained for several hours after eye-patch removal is surprising, because the effect of short-term MD normally decays within 90 minutes (see also Fig. S1)^24, 26^. Our results therefore indicate that sleep maintained visual homeostatic plasticity, stabilizing the ocular dominance change induced by MD and delaying the expected decay until the awakening. Interestingly, the effect of MD (deprivation index) measured before and after 2h of sleep did not correlate across subjects (Spearman’s rho=0.18, p=0.47). This suggests that individual sleep pattern could interact with visual homeostatic plasticity and that the instatement and maintenance of plasticity might be mediated by different neural processes, possibly reflected in different features of NREM sleep.

To rule out the effect of total darkness exposure occurring during sleep, we performed an additional condition during which participants, after MD, laid down in a completely dark room for 2 hours, without sleeping (Monocular Deprivation Morning, MDmorn, Figure 2A). The experiment was performed in the morning, to prevent the occurrence of sleep during the two hours of dark exposure. In this experiment (Figure 2B, blue symbols), we found that the effect of MD (mean DI ± SE = 0.71 ± 0.05, two-tailed, one-sample t-test, t(16)=-4.91, p<0.0002, Cohen’s d = 1.19) decayed to baseline within 2 hours of darkness (mean DI ± SE = 1.03 ± 0.07, two-tailed, one-sample t-test, t(16) = -0.04, p = 0.96, Cohen’s d = 0.009), similarly to what observed with normal visual stimulation after patch removal. We then directly compared the decay of the effect of deprivation after two hours of sleep (Figure 2B, red symbols) or after two hours of dark exposure (Figure 2B, blue symbols) by performing a 2 (TIME, before and after) x 2 (CONDITION, MDnight and MDmorn) repeated measures ANOVA. We found a significant interaction between the factors CONDITION and TIME (repeated measures ANOVA, F(1,16) = 4.48, p = 0.05, η2 = 0.22), confirming the specific role of sleep in stabilizing visual homeostatic plasticity induced by monocular deprivation.

**Figure 2.**
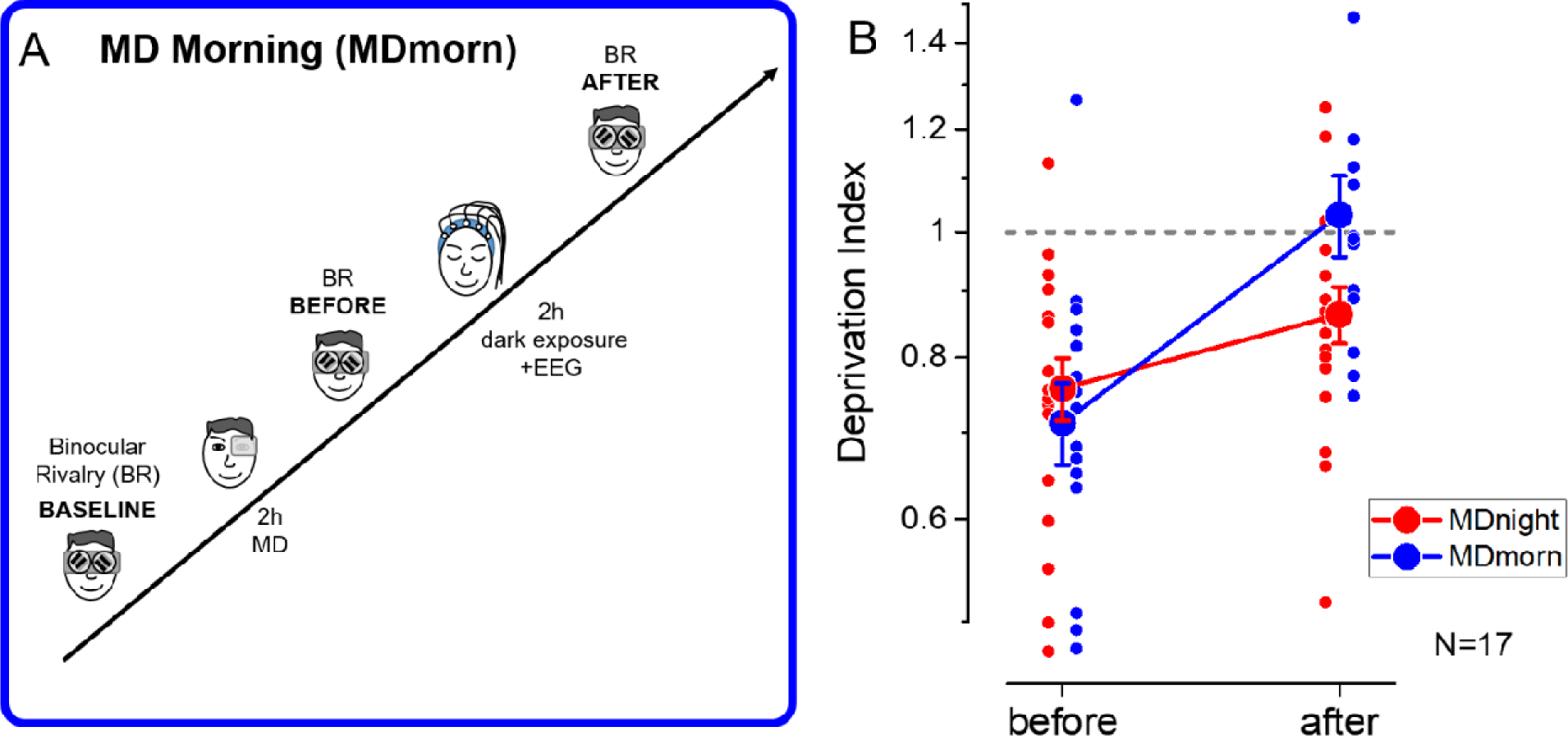
Experimental paradigm and results of the dark exposure (Monocular Deprivation Morning, MDmorn) condition. A) Diagram of the experimental paradigm for the MD morning condition. After 2h of MD, participants spent 2h in total darkness, while their EEG activity was recorded. Following the 2h of dark exposure, ocular dominance was assessed by binocular rivalry. (B) The deprivation index measured before and after 2h of dark exposure without sleep performed in the monocular deprivation morning session (blue symbols) and before and after 2h of sleep in the MD night session (red symbols), for the 17 participants who performed both conditions. Error bars represent 1±SEM, the small dots represent individual subjects, the big dots represent the average.

The two hours of sleep and dark exposure took place at different times of the day, one late at night and the other one early in the morning. To rule out a possible influence of the circadian rhythm on homeostatic plasticity and its decay, we performed a control experiment in which we measured the effect of 2h of monocular deprivation early in the morning or late at night in a separate group of adult volunteers. We found that the dynamics of the monocular deprivation effect were similar for the morning and evening sessions, as in both cases ocular dominance returned to baseline levels within 120 minutes (repeated measures ANOVA, TIME*CONDITION: F(4,32) = 1.08, p = 0.38, η2 = 0.12). Moreover, the MD effect was significantly larger when deprivation was performed in the morning (Figure S1, repeated-measures ANOVA, CONDITION: F(1,8)=6.87, p=0.031, η2=0.46), indicating a lower plastic potential of the visual cortex in the evening. Taken together, these results indicate that the maintenance of the MD effect is specific to sleep. Moreover, the lower plastic potential observed in the evening indicates that the role of NREM sleep in stabilizing homeostatic ocular dominance plasticity might be underestimated.

Having demonstrate a stabilization of the boost of the deprived eye with sleep, two different but intermingled issues arise: 1) how MD affects subsequent sleep, and 2) how sleep contributes to stabilize visual homeostatic plasticity. As the first 2 hours of night sleep contained none or just a few minutes of REM sleep, both effects have been investigated within NREM sleep.

NREM sleep features were derived from the 2 hours of EEG recording before the first night awakening. These include the power scalp distribution of slow wave activity (0.5-4Hz) and sigma (9-15Hz) rhythms, the rate (waves per time unit) and shape of SSO as well as the density (waves per time unit) and power of sleep spindles.

Table S1 shows descriptive statistics of sleep architecture parameters in the MDnight and Cnight. Also, it provides between-nights statistical comparisons and the study of putative association between sleep architecture, susceptibility to MD (DI before) and stability of the effect during sleep (DI after). No sleep architecture parameters varied significantly between nights or were associated with DI measurements.

Given previous evidence of ocular dominance homeostatic plasticity at level of early visual cortex^30^, we analysed the EEG rhythms and patterns in an extended occipital ROI reflecting the activity of the majority of primary and associative visual areas. We compared the changes over the ROI from the control to experimental night of each feature. A control ROI was selected in correspondence of the sensory-motor cortex (Figure S2).

Large ROIs have the advantage to compensate for the individual large variations of source dipole localization across visual areas and to decrease uncorrelated neuronal noise by averaging. We also performed single electrode analysis obtaining similar, although noisier, results (see scalp maps in Figure 3 and 4, and single electrode correlations distributions in Figure S3 and S4). We never observed any main changes of the sleep rhythms or features between MD and Control night recordings, in any of the ROIs (Table S2). However, we observed a strong correlation between changes in sleep features and ocular dominance plasticity measured before sleep in the occipital ROI (Figure 3 and Figure S3 for the coherence of correlations between electrodes within ROIs).

**Fig 3.**
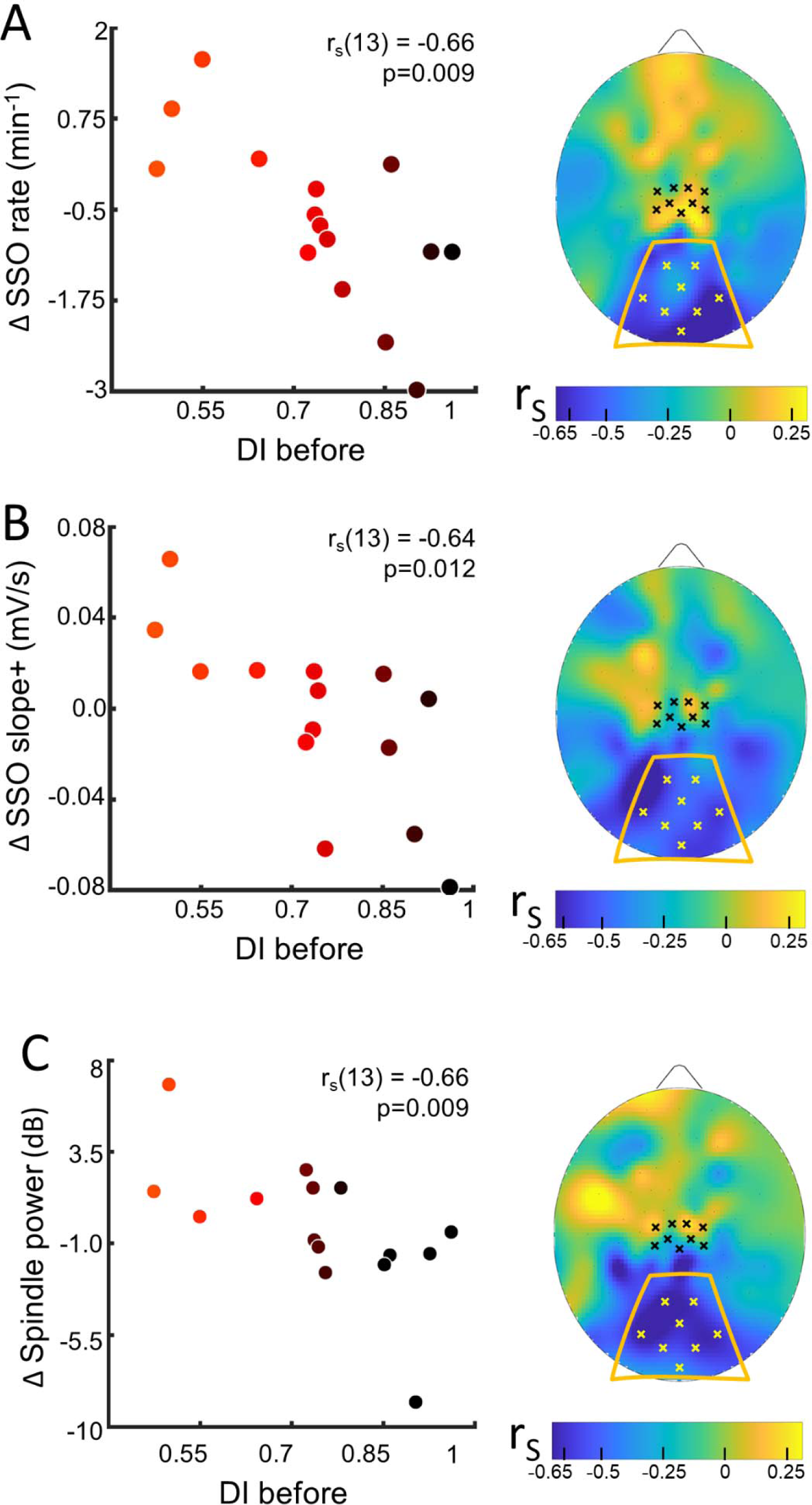
Sleep slow oscillation and spindle modulation with the eye dominance induced by monocular deprivation plasticity index. (A) The changes from Control to MD night of the rate of SSOs was correlated with the deprivation index measured before sleep (DI before). Scatterplot shows individual values averaged within the occipital ROI (Spearman’s r with the p-value shown as an inset in the scatterplot). Color of dots spans from black to red as a function of individual plasticity. No significant correlation appeared when considering the control ROI defined in the sensory-motor cortex. The scalp map shows the spatial distribution of correlation as estimated electrode by electrode; within the map, yellow and black dots mark electrodes belonging to the occipital and sensory-motor ROIs, respectively).(B) same as A for the steepness of slope+ of SSOs; (C) same as A for the spindle power

**Fig 4.**
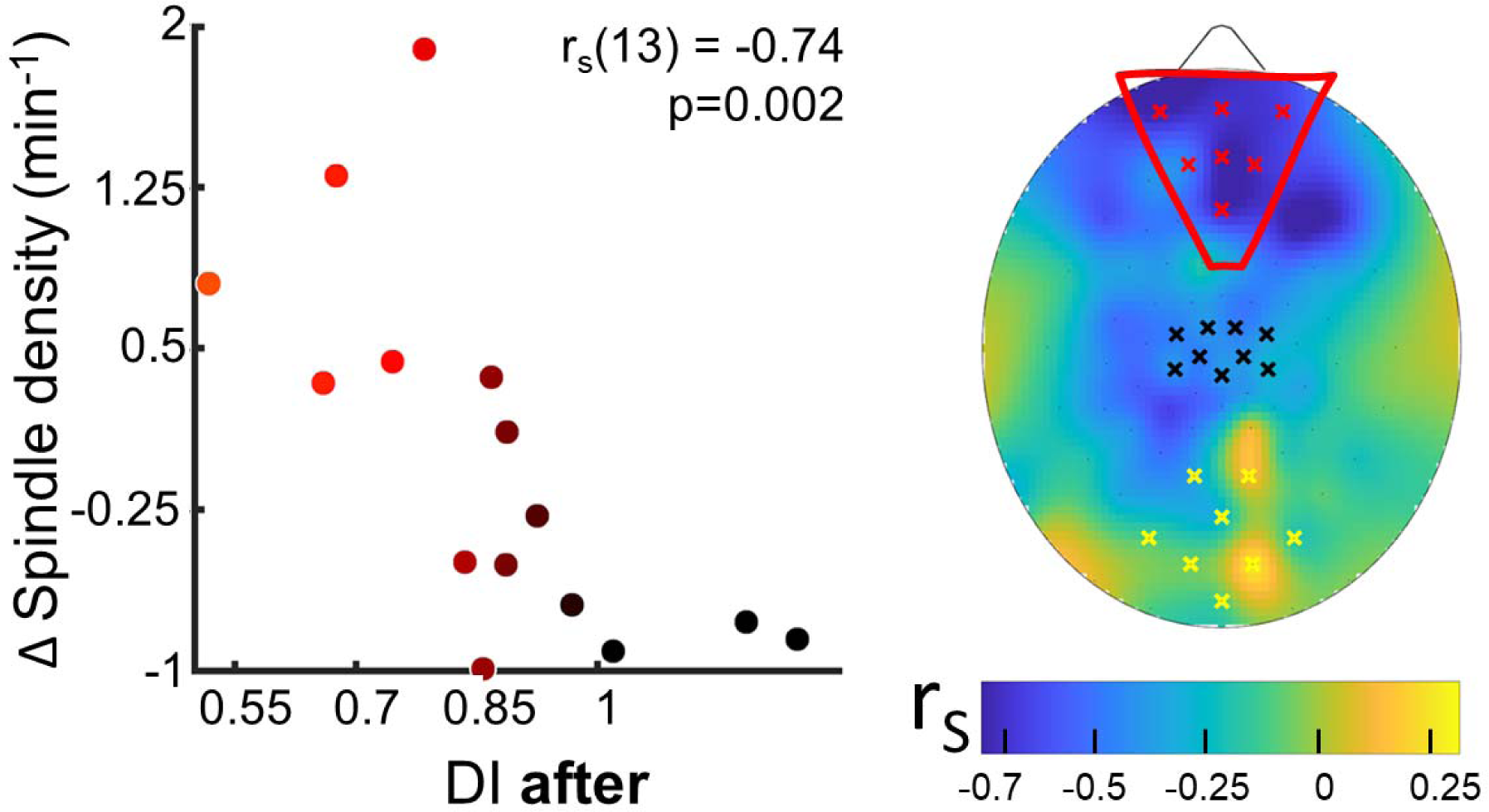
Spindle density modulation with the residual plasticity after two hours of sleep. The changes from Control to MD night of the spindle density were correlated with the deprivation index measured after sleep (DI after). Scatterplot shows individual values averaged within the prefrontal ROI (Spearman’s r with the p-value shown as an inset in the scatterplot). Color of dots spans from black to red as a function of individual DI after. No significant correlation appeared when considering ROIs defined in the occipital and in sensory-motor cortex. The scalp map shows the spatial distribution of correlation as estimated electrode by electrode over the scalp; within the map yellow, black and red dots mark electrodes belonging to the occipital, sensory-motor and prefrontal ROIs, respectively).

For the SSOs, a strong correlation with plasticity was observed between changes in SSO rate (rho=-0.66, p=0.009, p_fdr_=0.036, Figure 3, A) and shape (slope+, rho=-0.64, p=0.012, p_fdr_=0.037, Figure 3, B; SSO amplitude, rho = -0.56, p=0.03, p_fdr_=NS, Figure S4, A) at the level of the occipital ROI: subjects showing a stronger plasticity effect showed 1) increased SSO rate, 2) increased sharpness of transitions from the SSO negative peak, whereas subjects with lower plasticity effect exhibited opposite parameters changes.

The opposite changes between subjects with high and low plasticity, without any main effect of MD manipulation on SSO features, suggest different local synchronization in the sleeping neurons undergoing the bistability behaviour of SSO. Finally, for the sleep spindles, a correlation with plasticity was observed between their mean power (rho=-0.66, p=0.009, p_fdr_=0.036; Figure 3C): power of sleep spindles over the occipital sites increased in the MD night for participants showing higher visual homeostatic plasticity (i.e. a large boost of the deprived eye), while the power was reduced in participants showing a lower response to MD. This result, together with the other sigma power estimates, reinforces the idea behind a modulation of thalamocortical interaction as a homeostatic res to MD. Indeed, sigma activity power expressed during the whole NREM sleep in the occipital ROI correlated with individual ocular dominance shifts (rho=-0.66, p=0.009, p_fdr_=0.036, Figure S4, B) and overlapping result was observed when considering the sigma rhythm expressed just before SSO events that favour the emergence of full-fledged SSO (rho=-0.70, p=0.0046, p_fdr_=0.025, Figure S4, C). These strong correlations contrast with the absence of association with the power of slow wave activity during the whole NREM sleep with the plasticity index (Figure S4, D).

The observed significant correlations were coherent at the single electrode level within the occipital ROI (Figure S3), but they were not observed in the sensory-motor ROI or at the electrode level within this control ROI. Also they were specific for sleep as no correlation between EEG rhythms power and visual plasticity was observed in the control experiment (darkness exposure condition, Figure S5).

To investigate how sleep contributes to the maintenance of the acquired ocular dominance we also analysed sleep features in frontal electrodes. We correlated ocular dominance plasticity measured after night awakening (Figure 1A, *binocular rivalry after*) with sleep characteristics in the three target ROIs measured in the early night: the occipital, the control sensory-motor and a new prefrontal ROIs (Figure 4 and Figure S6 for the coherence of correlations across electrodes within each ROI).

Sleep spindles density in the prefrontal ROI strongly correlated (rho=-0.74, p=0.002, p_fdr_=0.04) with the effect of MD as maintained after sleep (Figure 4): participants retaining the stronger effect after 2h of sleep showed substantial increase of spindle density. No correlation was observed in the other ROIs. Besides, neither power of slow wave activity and sigma bands, nor SSO rate and shape, were associated with the residual eye dominance after two hours of sleep (Figure S7).

Finally, we considered the variations of EEG parameters during the two hours of sleep to verify the stability of the sleep features as a function of the delay from deprivation. Among the considered EEG measures (Figure S8), none was associated with the residual eye dominance after two hours of sleep (DI after).

## Discussion

We investigated the interplay between visual homeostatic plasticity and sleep in healthy adult humans after a short MD period. We report for the first time that different features of NREM sleep affect and are affected by homeostatic ocular dominance plasticity in adult humans: the plastic potential of the visual cortex is reflected by the expression of SSO and sigma activity in occipital areas and sleep consolidates the effect of short-term MD via increased spindles density in prefrontal area.

### Role of sleep in supporting visual plasticity

We found that sleep promotes the stabilization of visual homeostatic plasticity induced by 2 hours of MD: the deprived eye boost observed after MD, which normally decays within 90 minutes of wakefulness ^26, 27, 56^, is maintained by sleep for up to 6 hours after eye-patch removal. Importantly, this effect was specifically induced by sleep: two hours of dark exposure after MD did not prevent the decay of the effect. Finally, we found that the circadian rhythm did not influence the dynamics of the effect, as a similar time-course of plasticity was observed when MD was performed early in the morning or late at night.

Results from animal models have shown that NREM sleep consolidates ocular dominance plasticity during the critical period^36–38^. This sleep-dependent consolidation relies on Hebbian mechanisms and it is mediated by synaptic potentiation through increased NMDAR and PKA activity^36^ and decreased GABAergic inhibition^38^, potentiating the input that more successfully drives the output activity. This type of Hebbian plasticity has never been observed in ocular dominance plasticity in adult humans, while it has consistently reported for homeostatic plasticity, the capacity to upregulate the gain of the weak input. Here we show for the first time that NREM sleep has an effect on ocular dominance plasticity also past the critical period in humans.

The sleep homeostatic hypothesis ^57^ should not be confused with the homeostatic process that drives this type of plasticity. For OD the homeostatic mechanism acts on the overall activity both for the deprived and non-deprived eye, while the sleep homeostatic hypothesis states that during NREM sleep, synaptic weights only down-scale in compensation for neural activity during wakefulness^57^. While the theory is validated by a large set of experimental findings, it may not be easily directly linked to our results.

The homeostatic plasticity process triggered by MD is associated with up- regulation of the deprived eye activity, but also with down- regulation of the non-deprived eye at the level of the primary visual cortex.^28–30, 43^ NREM sleep appears therefore to intervene in a rebalancing process involving overall visual cortical activity. This rebalancing process operates in opposite directions for the deprived and non-deprived eye, which explains the lack of an association between sleep features at the occipital level and the maintenance of the MD effect observed after sleep as well as the lack of a main effect of MD on the subsequent sleep features.

Interestingly, we found that the influence of sleep on ocular dominance plasticity was reflected by some sleep features in high-level areas, indicating a role of extra-striate cortex in regulating visual plasticity in adults. The density of sleep spindles was associated with the MD effect maintained after sleep in a distinct cluster of electrodes distributed over the mesial frontal and prefrontal cortex. Sleep spindles have been described as replay events of new information acquired during wakefulness and in the course of consolidation during NREM sleep, which might mark phasic activations of a circuit involving the hippocampus^58^: indeed the consolidation of several forms of plasticity has been associated with sleep spindle density^59, 60^ and some forms of visual plasticity (e.g. perceptual learning) are reflected in changed activity and connectivity within the hippocampus^61^, which has dense anatomical connections with the primary visual cortex^62^. This is a surprising results given the large evidence that the mechanisms underlying ocular dominance are in primary visual cortex. However, we cannot exclude that the primary visual cortex may be under hippocampal control in wakefulness, as visual memory may require replay activity. Interestingly simple orientation texture discrimination tasks are also stabilized during sleep and also in these case frontal activity are implicated in the stabilization process^51^.

Altogether, consistent with other results, we show that activity in non-visual areas plays a role in modulating the decay of short-term visual plasticity in adult humans and that this activity might be crucial to promote the stabilization of the plastic changes induced by MD that we observed in amblyopic patients^33^. In amblyopia, the boosting effect after MD was stabilized across consecutive days with sleep occurring in between sessions and became permanent after performing short-term MD over 4 weeks.

### Sleep and the plastic potential of visual cortex

We found that the expression of SSO in visual areas reflects the interindividual variability in visual homeostatic plasticity: their rate increased and their shape changed in occipital sites proportionally to the shift in ocular dominance induced by MD, as measured immediately before sleep.

Less SSOs observed in low plasticity subjects also means less downstate periods. During the downstate a large majority of neurons are silent for fractions of second and restorative processes occurring at the level of individual brain cells occur^63^. Thus, less SSOs could indicate an average lower need of such processes due to the previous sensory deprivation. In this line, also healthy volunteers who sleep after blindfolding exhibit a dramatic decrease of SSOs^64^. At the other end, more SSOs in subjects with high visual plasticity could indicate the homeostatic activation of this mechanism endorsing the ocular dominance shift. SSO shape changed accordingly after MD: subjects with high visual plasticity show greater SSOs with steeper downstate exit slope (slope+) compared to their basal characteristics. Larger SSOs indicate larger groups of cortical neurons synchronously involved in these bistable events, while a steeper positive slope has been associated with a stronger coupling with thalamic structures ^65, 66^ .

Altered activity in the slow wave frequency band (the band including SSOs) was also observed in sleep subsequent sensorimotor deprivation^67^, however SSO events were not studied directly in this experimental model. The type of plasticity induced by sensorimotor deprivation differs from the effect of short-term MD, consistently with Hebbian mechanisms mediating sensory-motor plasticity and homeostatic mechanisms mediating short-term ocular dominance plasticity. Short-term MD induces opposite changes in VEPs, increasing VEPs amplitude of the deprived eye and decreasing that of the non-deprived eye^67^, while sensorimotor deprivation reduces SEPs/MEPs related to the immobilized arm leaving unaltered SEPs/MEPs related to the free arm (Huber et al., 2005). These differences might explain the different pattern of sleep changes observed after sensorimotor deprivation and after MD: while Huber et al (2006)^67^ reported a significant decrease in slow wave activity power, we did not observe a main effect of our experimental manipulation, reflecting balanced overall activity changes of the deprived and non-deprived eye between the pre- and post- deprivation state. However, we reported a strong correlation between SSO rate/shape parameters and the MD effect, suggesting that SSO encodes homeostatic plasticity in the adult visual cortex.

The expression of oscillating activity in the sigma band also changed as a function of the individual plastic potential of the visual cortex. Most of the sigma activity during NREM sleep relies on the pacemaker cells within the reticular thalamic nucleus forcing patterned activity in thalamocortical and cortical cells. The paradigmatic expression of this activity is the sleep spindle during which ideal conditions for fine scale plasticity occur: thalamic inputs during spindles yield to dendritic depolarization but keeping the cell from firing and triggering calcium entry into dendrites^68^. Recordings in vivo from lateral geniculate nucleus and visual cortex have shown that SSOs and sigma activity are strongly coordinated within thalamocortical circuits^69^, and evidence are accumulating for reticular thalamic plasticity sustaining spindle-based neocortical– hippocampal communication^17^. Therefore, during NREM sleep, any factor modulating the activity of one of the two structures affects both and this lays the foundation for coordinated cortico-thalamic plastic changes^4, 65, 70–72^.

Overall, these results suggest that SSOs and sigma activity reflect the degree of homeostatic plasticity induced by short-term MD. Both homeostatic plasticity^45, 73^ and SSO^47, 74^ are linked to GABAergic inhibition, the observed effect could therefore in principle be mediated by a change in excitation/inhibition balance in the visual system. Importantly, in humans, GABA concentration measured after 2 hours of MD in V1 decreased proportionally to the observed ocular dominance shift^32^, and slow wave activity expression increases in response to GABA agonists administration^75^. Interestingly, a recent study^76^ reported that complementary changes in the excitation/inhibition balance measured in the visual cortex of adult humans during NREM and REM sleep are correlated with visual plasticity induced by perceptual learning, further pointing to a leading role of GABAergic inhibition in mediating these two phaenomena. We speculate that the alteration of SSO shape and density observed at the occipital level during the sleep immediately following monocular deprivation might reflect the change in GABA concentration induced by deprivation and be a neurophysiological marker of the interindividual variability in the level of plasticity in adult humans.

## Conclusions

Sleep oscillatory activity can reflect the plastic potential of the occipital cortex: people highly susceptible to a visual manipulation (short-term monocular deprivation) affecting primary visual cortex activity show increased SSO and sigma activity in occipital sites. Sleep can also extend for many hours an otherwise transient unbalance of visual cortical activity, and sleep spindles in prefrontal regions appear to support the process as subjects exhibiting stronger maintenance had increased frontopolar sleep spindle density, as it occurs for many memory consolidation processes.

## Materials and methods

### Participants

Nineteen healthy volunteers (mean age ± SD 24.8 ± 3.7 years, range 21-33 years; 8 males), participated in the main study. The eligibility of each volunteer was verified by semi-structured interviews conducted by a senior physician and psychiatrist (AG) based on the following inclusion criteria: no history of psychiatric/neurological disorders (including sleep disorders), being drug free for at least one month. All subjects had normal or corrected-to-normal visual acuity (ETDRS charts). Enrolled volunteers received the following instructions to be accomplished in each day of the experimental procedures: to avoid any alcohol intake, coffee intake in the evening before sleep sessions and physical work out just before all the experimental sessions (both the night and the morning sessions). All subjects performed the three experimental conditions, except for one subject, who did not perform the control night condition, and two subjects who did not perform the monocular deprivation morning condition, because of personal problems. Four participants were excluded from the EEG analysis because poor signal quality in the EEG signal occurred during the sleep recording.

Sample size was determined based on previous studies on sleep and plasticity (e.g. visual^76–78^ and visuo-motor learning^12, 79, 80^) using a within-subject design, indicating that robust results can be observed with a sample size of 10-15 participants. In addition, since this is the first study exploring the role of sleep in homeostatic plasticity induced by MD and that for its nature it has to take into account the complexity of sleep as expressed through different parameters, this sample size and the need for multiple tests corrections have led to highlight only effects characterised by strong effect size.

Nine additional volunteers (mean age ± SD 25.7 ± 5.2 years, range 20-37 years; 3 males) participated in the control experiment. all participants had normal or corrected-to-normal visual acuity (ETDRS charts) and no history of psychiatric/neurological disorders.

### Ethics statement

All eligible volunteers signed an informed written consent. The main study was performed at the University of Pisa (Italy). The experimental protocol was approved by the Local Ethical Committee (Comitato Etico Pediatrico Regionale—Azienda Ospedaliero-Universitaria Meyer— Firenze), under the protocol “Plasticità del sistema visivo” (3/2011) and complied the tenets of the Declaration of Helsinki (DoH-Oct2008). Participants were reimbursed for the time at a rate of 100€ per night and 50€ for the morning session experiment. The control experiment was performed at the Laboratoire des Systèmes Perceptifs in Paris (France). The experimental protocol was approved by the local ethics committee (Comité d’éthique de la Recherche de l’Université Paris Descartes, CER-PD:2019-16-LUNGHI) and was performed in accordance with the Declaration of Helsinki (DoH-Oct2008). All participants gave written informed consent. The naïve participants received financial compensation of 10€ per hour.

### Monocular deprivation

Monocular deprivation (MD) was performed by applying a custom-made eye-patch on the dominant eye. Eye dominance was defined according to the binocular rivalry measurement performed in the baseline (11 right eyes were deprived in the main experiment and 5 right eyes in the control experiment). The eye-patch was made of a translucent plastic material that allows light to reach the retina (attenuation 15%) but completely prevents pattern vision. MD lasted 2h, during which participants stayed in the laboratory control room under experimenters’ supervision and did activities such as reading and working on the computer. In the last half hour all subjects underwent the EEG montage so that at the patch removal, after the acquired eye dominance measurement, they could go to sleep without any delay.

### Binocular Rivalry

Main study: visual stimuli were generated by the ViSaGe stimulus generator (CRS, Cambridge Research Systems), housed in a PC (Dell) and controlled by Matlab programs. Visual stimuli were two Gaussian-vignetted sinusoidal gratings (Gabor Patches), oriented either 45° clockwise or counterclockwise (size: 2s = 2°, spatial frequency: 2 cpd, contrast: 50%) displayed on a linearized 20inch Clinton Monoray (Richardson Electronics Ltd., LaFox, IL) monochrome monitor, driven at a resolution of 1024×600 pixels, with a refresh rate of 120 Hz. To facilitate dichoptic fusion stimuli were presented on a uniform grey background (luminance: 37.4 cd/m2, C.I.E: 0.442 0.537) in central vision with a central black fixation point and a common squared frame. Subjects received the visual stimuli sitting at the distance of 57 cm from the display through CRS Ferro-Magnetic shutter goggles that occluded alternately one of the two eyes each frame.

Each binocular rivalry experimental block lasted 3 minutes. For each block, after an acoustic signal (beep), the binocular rivalry stimuli appeared. Subjects reported their perception (clockwise, counter clockwise or mixed) by continuously pressing with the right hand one of three keys (left, right and down arrows) of the computer keyboard. At each experimental block, the orientation associated to each eye was randomly varied so that neither subject nor experimenter knew which stimulus was associated with which eye until the end of the session, when it was verified visually.

Control experiment: The visual stimuli were generated in Matlab (R2020b, The MathWorks Inc., Natick, MA) using Psychtoolbox-3 (Brainard, 1997) running on a PC (Alienware Aurora R8, Alienware Corporation, Miami, Florida, USA) and a NVIDIA graphics card (GeForce RTX2080, Nvidia Corporation, Santa Clara, California, USA). Visual stimuli were presented dichoptically through a custom-built mirror stereoscope and each subject’s head was stabilised with a forehead and chin rest positioned 57 cm from the screen. Visual stimuli were two sinusoidal gratings oriented either 45° clockwise or counter clockwise (size: 2°, spatial frequency: 2 cpd, contrast: 50%), presented on a uniform grey background (luminance: 148 cd/m2, CIE x=.294, y=.316) in central vision with a central white fixation point and a common squared white frame to facilitate dichoptic fusion. The stimuli were displayed on an LCD monitor (BenQ XL2420Z 1920 x 1080 pixels, 144 Hz refresh rate, Tapei, Taiwan). Each binocular rivalry experimental block lasted 3 minutes. For each block, after an acoustic signal (beep), the binocular rivalry stimuli appeared. Subjects reported their perception (clockwise, counterclockwise or mixed) by continuously pressing with the right hand one of three keys (left, right and down arrows) of the computer keyboard. At each experimental block, the orientation associated with each eye was randomly varied.

### High density EEG recordings

EEG was recorded using a Net Amps 300 system (Electrical Geodesic Inc., Eugene, OR, USA) with a 128-electrodes HydroCel Geodesic Sensor Net. The EEG system employs a high spatial density electrode system with full head coverage including periocular, cheekbone and neck sensors. This allows the detection of both vertical and horizontal eye movements and muscle tone (both from the zygomaticus major muscles and upper trapezius muscles on the neck) directly from the EEG cap. During the two hours of recordings, electrode impedances were kept below 50 KΩ and signals were acquired with a sampling rate of 500 Hz, using the Electrical Geodesic Net Station software, Version 4.4.2. EEG recordings were analysed using tailored codes written in Matlab (MathWorks, Natick, MA, USA) and EEGLAB toolbox functions^81^.

### Experimental procedures

#### Main Study

Experimental procedure comprises three sessions, and for each volunteer, sessions were completed within a month and at least one week apart. The experiment took place in a dark and quiet room, with a comfortable bed equipped for EEG recordings and the apparatus for measuring binocular rivalry placed next to the bed. Each volunteer spent two nights (from 9:30 PM to 8AM) and a morning (from 9 AM to 2 AM) at the laboratory: i) a Monocular Deprivation Night (MDnight), in which participants underwent 2h of MD before sleep; ii) a Control Night (Cnight), in which no MD was performed before sleep, but participants waited two hours in the laboratory performing the same activities and undergoing eye dominance measures at the same times as in MDnight; iii) a Monocular Deprivation Morning (MDmorn), during which subjects, after the MD, lied in the same bed of night sessions, resting in the dark for two hours, avoiding sleeping, in order to study the acquired eye dominance extinction without visual stimuli. The order of the night sessions was randomized and balanced within the experimental group.

For the night sessions (MDnight and Cnight) the volunteer was welcomed in the laboratory around 9.30 PM and then tested for the baseline measurement of BR. Consequently, in the MDnight session around 10 PM the monocular patch was applied, the volunteer spent two hours reading or watching movies and thus, after exactly two hours, the patch was removed. The volunteer, already wearing the EEG cap, was tested again for the second measure of BR. To calculate the deprivation index (DI, see eq. 1) before sleep we used the first and second measure of BR. Once the second BR measurement was carried out, the subject was immediately invited to go to bed to fall asleep, this happened around midnight for both MDnight and Cnight. During both nights, sleep was interrupted after two hours (around 2.30 AM) to test the volunteer again for the third measure of BR (it took about 10 min), after which volunteers could sleep undisturbed until 7.30 AM (from 4 to 5 hours).

For assessing ocular dominance during night sessions, binocular rivalry was measured at four different times: before MD (or waiting for the Cnight, night *baseline*, 2×3min blocks), after 2h of MD and before sleep (or waiting for the Cnight, *before sleep*, 2×3min blocks), after the first 2h of sleep (*after sleep*, 2×3min blocks) and after the second awakening (*morning awakening*, 5×3min blocks measured 0, 5, 10, 15 and 30 min after eye-patch removal). Similarly, during the MDmorn session, binocular rivalry was measured at three times: before MD (morning *baseline*, 2×3min blocks), after 2h of MD and before dark exposure (*before dark*, 2×3min blocks), after the 2h dark exposure (*after dark*, 2×3min blocks).

During the night sessions, EEG was acquired from the in-bed time until subjects were woken up for performing the binocular rivalry measures after two hours of sleep, whereas during the MDmorn session, EEG was acquired in the two hours of dark exposure. According to EEG recordings, in the night sessions participants fell asleep easily and fast (sleep latency 9±2 min – mean±se), showed a normal organization of sleep structure (37±3% spent in N2 stage, and 50±3% spent in N3 stage, on average), exhibited none (8 out of 15 subjects in the MDnight and 4 out of 15 subjects in the Cnight) or few minutes of REM sleep (9±2 min) and none of them had wakefulness episodes after sleep onset.

For the MDmorn session, the subject was welcomed in the laboratory around 9 AM, shortly thereafter he/she was tested for the first BR measurement and then subjected to MD for two hours, during which he/she could read or watch movies. Consequently around 11.30 the patch was removed, the volunteer was tested for a second BR measurement immediately followed by two hours of resting by lying in bed in the dark. After 2 hours (1.30 PM), the BR was again measured multiple times according to a sequence suitable for reconstructing the extinction curve of the MD effect (the sequence lasted 30 minutes). In the MDmorn session none of the subjects were allowed to feel asleep as EEG was monitored in real-time and, in case of drowsiness signs (EEG slowing to theta), a bell tolled in the resting room. Having done this session in the morning, the use of the bell was extremely occasional: some subjects never showed signs of falling asleep, others were alerted at most 3 times with the bell in the last part of the resting state. A diagram of the experimental paradigm for the MDnight is reported in Figure 1A.

### Monocular deprivation and control nights EEG processing

To prepare EEG signals for the planned analysis, they underwent some preprocessing. EEG pre- processing and analyses were performed using tailored codes written in Matlab (MathWorks, Natick, MA, USA); scalp maps were obtained using EEGLAB Toolbox functions^81^.

For night sessions, scalp EEG signals were re-referenced to the average mastoid and scored according to the AASM criteria^80^. In particular, artefacts related to movements or muscle twitches were detected during the signal inspection performed for visual sleep stage scoring. EEG segments affected by artefacts were thus excluded from the analysis. Moreover, temporary or permanent decline in individual channel signal quality (often due to instability or loss of contact with the scalp during recordings) was studied on the basis of the signal statistical moments (EEGLAB Toolbox^77^). Channels characterized by outliers in the statistical moments were classified as ’bad’ channels and excluded from analysis. At the end of these pre-processing steps, all recordings have shown less than 10% of artefact-contaminated segments.

Sleep macrostructure was evaluated by extracting a set of time-domain parameters from the sleep staging annotations: sleep latency (time length of the transition from lights-off to the first N2 sleep episode, min); wake episodes after sleep onset (WASO duration, min); shift phase (sleep stage shift per time unit), sleep fragmentation (awakenings and shifts to lighter sleep stages per time unit), N2, N3 and rapid eye movement (REM) stages duration (min); and REM latency (time from sleep onset to the first REM sleep episode, min). Moreover, according to the sleep stage scoring, artefact-free NREM (N2 and N3) segments were analysed for estimating power band content and identifying sleep patterns such as SSOs and sleep spindles.

For the EEG signals analysis, only segments classified as NREM (N2 and N3) and free of artefacts were considered. For power band content of NREM sleep, two frequency bands of interest were considered: slow wave activity (SWA: 0.5–4 Hz), and sigma (σ: 9–15 Hz). Power densities were estimated applying a Hamming-windowed FFT on 10 sec consecutive EEG segments and log-transformed (dB). For each segment, electrode and band, the absolute power was estimated by averaging over its frequency bins. The absolute power of each band and electrode was then obtained averaging among segments.

### Monocular deprivation morning EEG processing

<u>For the MDmorn session, EEG was pre-processed fo</u>r managing diffuse artefacts with reconstruction of virtually clean traces. Thus, EEG signals were high pass filtered at 0.1 Hz (Chebyshev II filter) and notch filtered at 50 Hz and its first harmonic (100 Hz). Channels located on the forehead and cheeks which mostly contribute to movement-related noise were discarded, thus retaining thus 107 channels out of 128^82^. Epochs with signals exceeding 100μV were automatically discarded; retained signals were visually inspected for the removal of artefacts and noisy channels. Rejected signals were substituted with signals obtained via spline-interpolation^83^ and furtherly submitted to the Independent Component Analysis for separating and removing components expressing eye movements, heart beats, line and channel noise - component selection also supported by the AI system of ICLabel^84^. After artefact removal procedures, EEG signals were re-referenced to the average of the mastoid potentials^82, 85^.

The artifact-free EEG segments analysed for estimating power band content in five frequency bands of interest were considered: theta (4–8 Hz), alpha (8–12 Hz), low beta (12–20 Hz), high beta (20–30 Hz) and gamma (30-45 Hz). Power densities were estimated by applying a Hamming-windowed FFT on 4 sec consecutive EEG segments and log-transformed (dB). For each segment, electrode and band, the absolute power was estimated by averaging over its frequency bins. The absolute power of each band and electrode was then obtained averaging among segments.

### Sleep Slow Oscillation detection and characterization

SSO events within NREM sleep periods were detected and characterized using a previously published and validated algorithm^82, 86^. In summary, the algorithm first identifies full-fledged SSOs: each wave should comprise (a) two zero crossings separated by 0.3–1.0 s, the first one having a negative slope; (b) a negative peak between the two zero crossings with a voltage less than −80 μV; (c) a negative-to-positive peak amplitude of at least 140 μV. An SSO event is defined when simultaneous SSOs (tolerance of up to 200ms delay between negative wave peaks) are recognised on multiple channels Then, detected SSO events are completed by clustering full-fledged SSOs with concurrent similar waves, even if sub-threshold detected on the other EEG channels. These detection criteria naturally include all K-complexes^87^ but can also complete detection with otherwise neglected small waves.

For each subject, night session and electrode channel, SSOs were characterized by the following parameters: detection density (number of waves per minute), negative to positive peak amplitude (NP amp), slope from the negative peak (slope+)^88^. Moreover, the sigma-activity (9-15Hz) expressed in the 1s window preceding each SSO was estimated as a known thalamocortical entrainment marker functioning as precursor of SSOs emergence^89^.

### Sleep Spindle detection and characterization

The sleep spindle recognition was carried out according to the approach proposed and validated by Ferrarelli et al. (2007)^90^, with some minor adaptations; the actual procedure is summarized below.

EEG data for all NREM sleep periods were band-pass filtered between 12 and 16 Hz (–20 dB at 11 and 17 Hz) and for each channel, the upper and lower envelopes of the filtered signal were derived. The spindle detection was based on the signal derived as the point-by-point distance between the upper and the lower envelopes (signal amplitude). Because signal amplitude varies between channels, for each NREM period and channel a value was estimated as the mean signal amplitude increased by twice its standard deviation. Thus, for each channel a threshold was defined as the weighted average over the values calculated for each period, with the weights corresponding to the period lengths. Sleep spindles were thus identified as the fluctuations in the amplitude signal exceeding two times the threshold. Based on the detected spindles, for each EEG channel, the measures characterizing each sleep recording were the density (spindle events per time unit) and the spindle power. The spindle power was derived as the average over the spindle events of the sigma band (12-16 Hz) power of each detected wave.

### Binocular Rivalry

The perceptual reports recorded through the computer keyboard were analysed using Matlab, the mean phase duration and the total time of perceptual dominance of the visual stimuli presented to each eye and mixed percepts were computed for each participant and each experimental block. The three-minute blocks acquired after the morning awakening were binned as follows: 0-8 min, 10-18 min, 30 min.

The effect of MD was quantified by computing a deprivation index (DI), The effect of MD was quantified by computing a deprivation index (DI), using the same formula in Lunghi, Emir et al (2015)^32^, which demonstrated that DI correlates with the change in GABA concentration in the visual cortex of adult participants. The DI summarises in one number the change in the ratio between deprived and non-deprived eye mean phase duration following MD relative to baseline measurements according to the following equation:

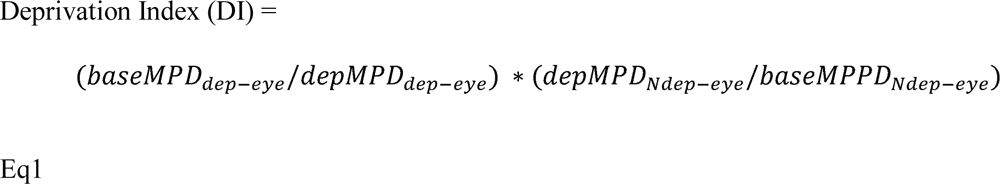

In Eq. 1, MPD is the Mean Phase Duration computed in seconds, base stands for baseline measurements, dep for measurements acquired after MD. A deprivation index value equal to 0 represents no change in the ratio between dominant and non-dominant eye mean phase duration, while a value >1 represents a decrease in dominant-eye predominance and a value <1 an increase in dominant eye predominance during binocular rivalry.

### Control Experiment

In order to investigate the influence of the circadian rhythm on visual homeostatic plasticity, the effect of monocular deprivation was tested for each participant in two different days, once in the morning, and once in the evening (the order of the conditions was counterbalanced across participants). In the morning session, the 2h monocular deprivation started at 9AM, while in the evening session, monocular deprivation started at 8PM. Each deprivation session was preceded by 2×3min baseline binocular rivalry blocks. After patch removal, we measured binocular rivalry continuously for 18 minutes in four separate 180sec-blocks, giving a short break every two minutes. Three-minute blocks of rivalry were tested again 30, 45, 60 90 and 120 minutes after restoration of normal binocular sight. The three-minute blocks acquired after monocular deprivation were binned as follows: 0-8 min, 10-18 min, 30-48 min, 60-93 min, and 120-123 min. Each participant therefore spent one morning (from 8.30 AM to 1 PM) and one evening (from 7.30 PM to 0.00 AM) in the lab. All the procedures and analyses were the same as the main study.

### Statistical analyses

The variation of deprivation indices measured during the two-night experiments was tested using repeated measures ANOVA with 1 factor (TIME) with 5 levels (*before, after, morning1, morning2, morning3*). In order to compare directly the decay of the deprivation effect in the MDnight and in the MDmorn condition, we used a repeated-measures ANOVA with 2 factors:

TIME (with 2 levels: before and after) and CONDITION (MDnight and MDmorn). For the control experiment, we used a repeated-measures ANOVA with 2 factors: TIME (with 5 levels corresponding to the five measurements obtained after deprivation, that is before, after, morning1, morning2, morning3) and CONDITION (MDnight and MDmorn). One-sample two-tailed t-tests tests were used for post-hoc tests, against the H0 mean=1. The Benjamini & Hochberg (1995)^91^ correction for multiple comparisons was applied for post-hoc tests, effect size was estimated computing Cohen’s d.

The variation of sleep characteristics according to the MD intervention and the dependence on the one hand on the acquired eye dominance (as measured by the deprivation index before sleep) and on the other on the residual dominance evaluated after two hours of sleep (as measured by the deprivation index after sleep) was evaluated considering three regions of interest (ROI): occipital ROI, prefrontal ROI and sensory-motor control ROI. These ROIs were selected on the literature evidence that MD alters activity over extensive occipito-parietal network of visual areas. The ROI in the prefrontal cortex was selected given its key role in organizing the ripple-mediated information transfer from hippocampus during NREM sleep ^92^. ROI-based analysis is not only in line with the assumptions of this work but also meets the low spatial resolution of EEG. Furthermore, employing ROIs allowed us to decrease the number of statistical tests (for each EEG feature, we have evaluated data on 3 ROIs instead of 90 electrodes). Finally, it made possible to manage inter-individual variability of anatomical brain structures. In particular, the large anatomical variability of V1 orientation implies a high variability in the dipole orientation and the related voltage over electrodes from Oz to CPz. The electrodes in the HydroCel Geodesic Sensor Net belonging to each ROI are shown in Figure S2.

For the effect of MD intervention on cortical activity during NREM sleep, the Wilcoxon signed rank test was applied on all the EEG sleep features. To this aim, each sleep feature was averaged over the electrodes belonging to occipital and sensorimotor control ROIs and thus compared between nights. Only variations between conditions whose statistical significance survived the false discovery rate (FDR^91^) correction for multiple testing were considered.

For the dependence on the individual plasticity induced by MD, the non-parametric Spearman correlation was determined between DI before sleep, and each EEG sleep feature averaged over the occipital and sensorimotor control ROIs. Only correlations whose statistical significance survived the FDR correction for multiple testing were considered. Analogously, the association with the individual residual eye dominance after sleep was determined by Spearman correlation calculated between DI after and each EEG sleep feature averaged over the occipital, the prefrontal and sensorimotor control ROIs. Also for this group of tests, only correlations whose statistical significance survived the false discovery rate (FDR88) correction were considered.

For all tests, the level of statistical significance after FDR correction was set at p<0.05; and the false discovery rate was set equal to p=0.05.

## Supporting Information

S1 Appendix includes supplementary tables and figures for Methods and Results.

## Supplementary Information Appendix

**Figure S1.**
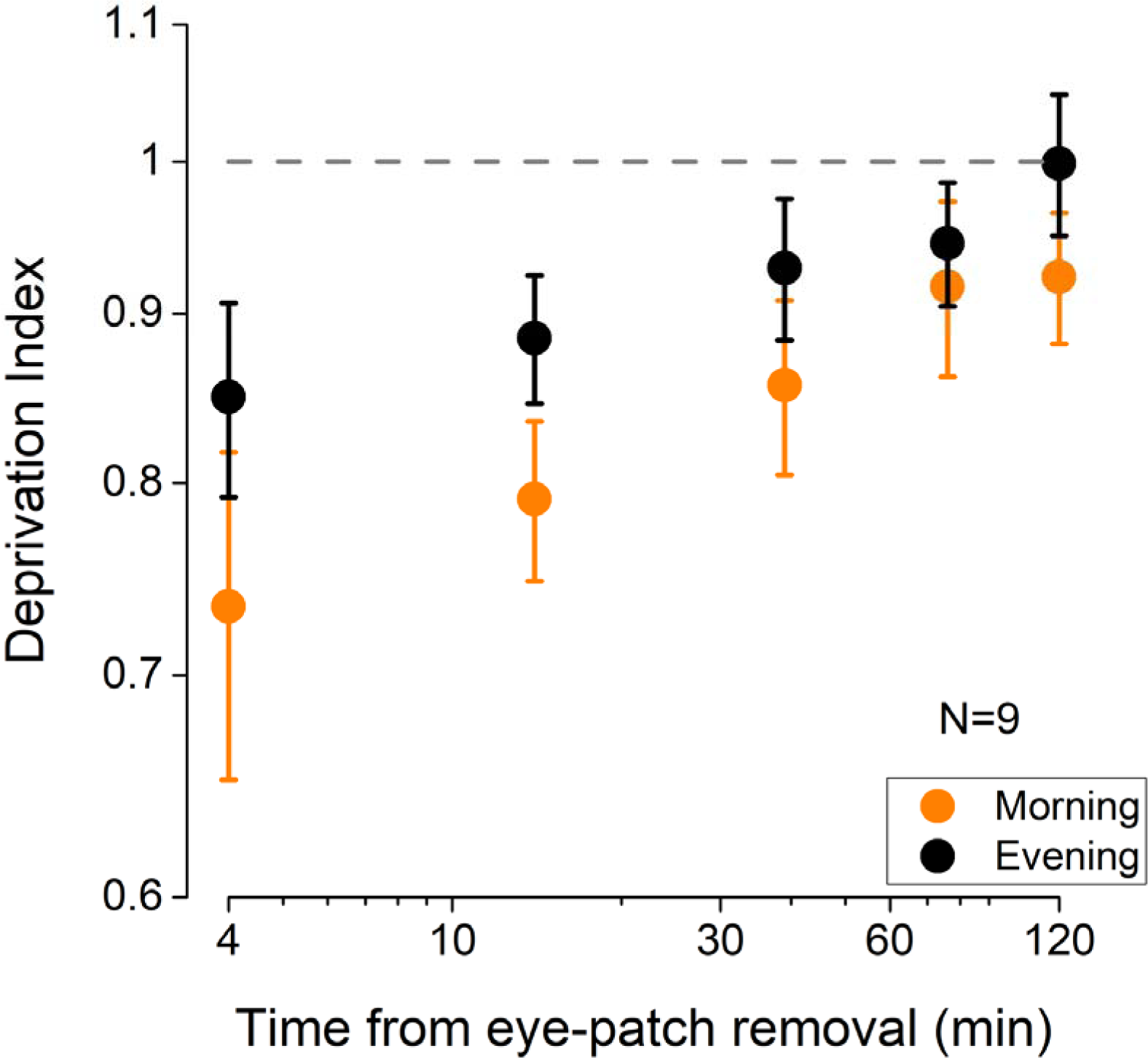
Control experiment results. The effect of 2h of monocular deprivation (Deprivation Index) on ocular dominance measured either early in the morning (orange symbols) or late in the evening (black symbols). The MD effect was significantly larger when deprivation was performed in the morning (repeated-measures ANOVA, CONDITION: F(1,8)=6.87, p=0.031, η2=0.46), indicating a lower plastic potential of the visual cortex in the evening.

**Figure S2.**
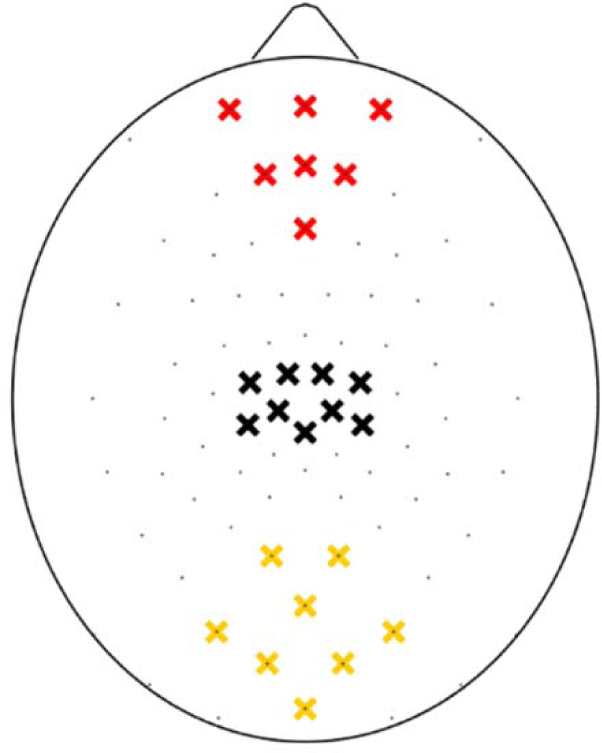
Electrodes in the HydroCel Geodesic Sensor Net belonging to each ROI. Yellow crosses mark electrodes belonging to the occipital ROI, red crosses mark electrodes belonging to the prefrontal ROI and black crosses mark electrodes belonging to the sensory-motor control ROI.

**Figure S3.**
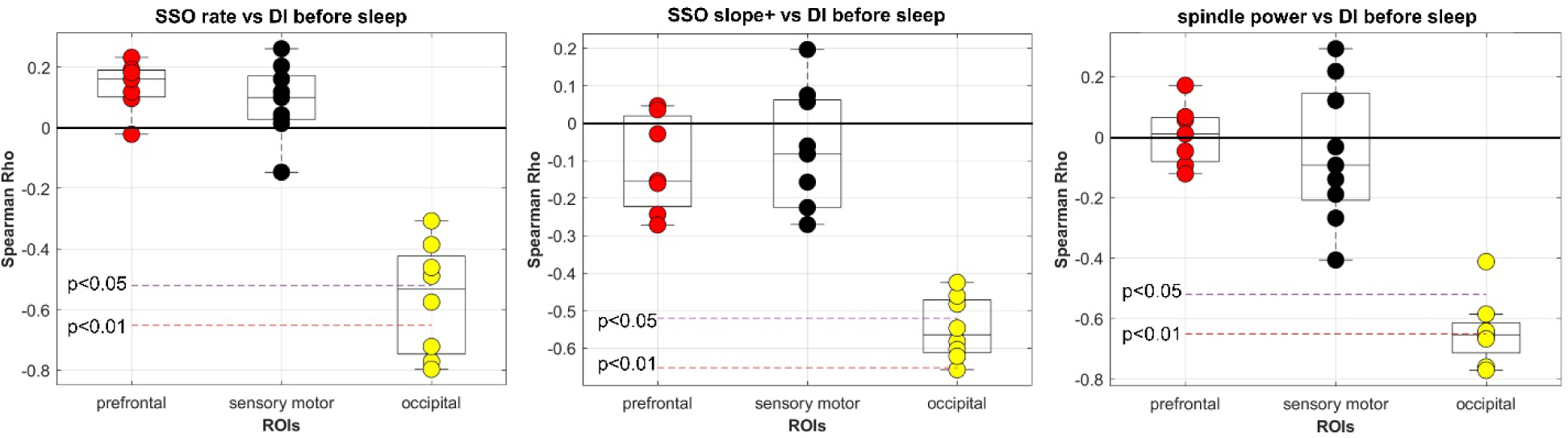
Correlations with DI before sleep. Coherence between electrodes within each ROIs for the features in Figure 3. Each dot corresponds to an electrode within the indicated ROI. Only electrodes belonging to the occipital ROI showed significant correlations with DI measured before sleep.

**Figure S4.**
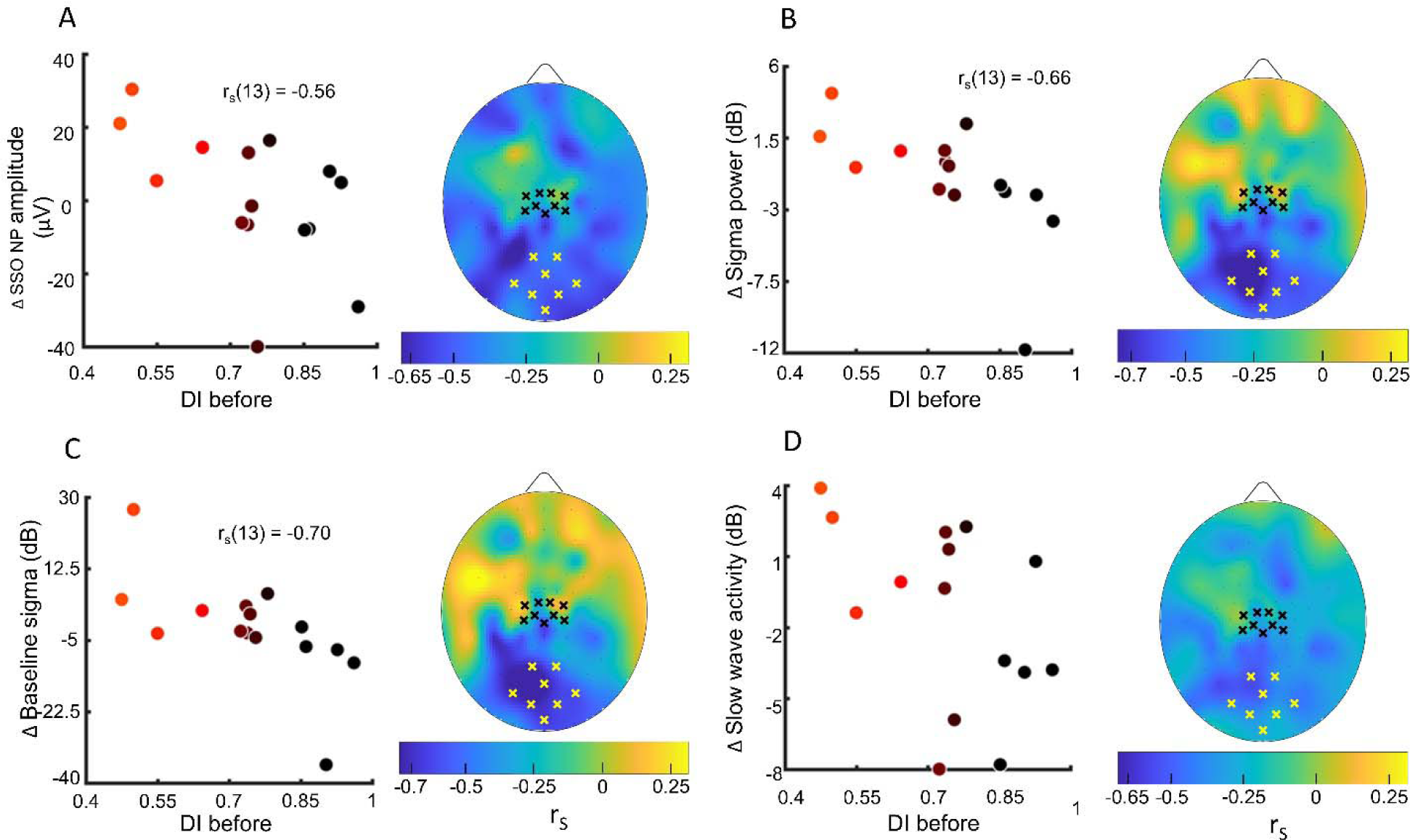
A) Correlation between the deprivation index measured **before** sleep (DI before) and the changes from the control night to the MD night of the mean sleep slow oscillations (SSO) amplitude measured . Scatterplot of the SSOs amplitude averaged over the occipital ROI (yellow crosses mark electrodes belonging to the ROI) with the deprivation index. Dot’s colour changes from black to red as a function of DI. The control ROI defined in the sensory-motor cortex (black crosses) shows no significant correlation with DI. The scalp map shows the spatial distribution of correlation; (B) same as A for the sigma activity power; (C) same as A for the baseline sigma rhythm expressed before SSO events; (D) same as A for the power of slow wave activity. For (D) no significant correlation was observed when averaging over both ROIs.

**Figure S5.**
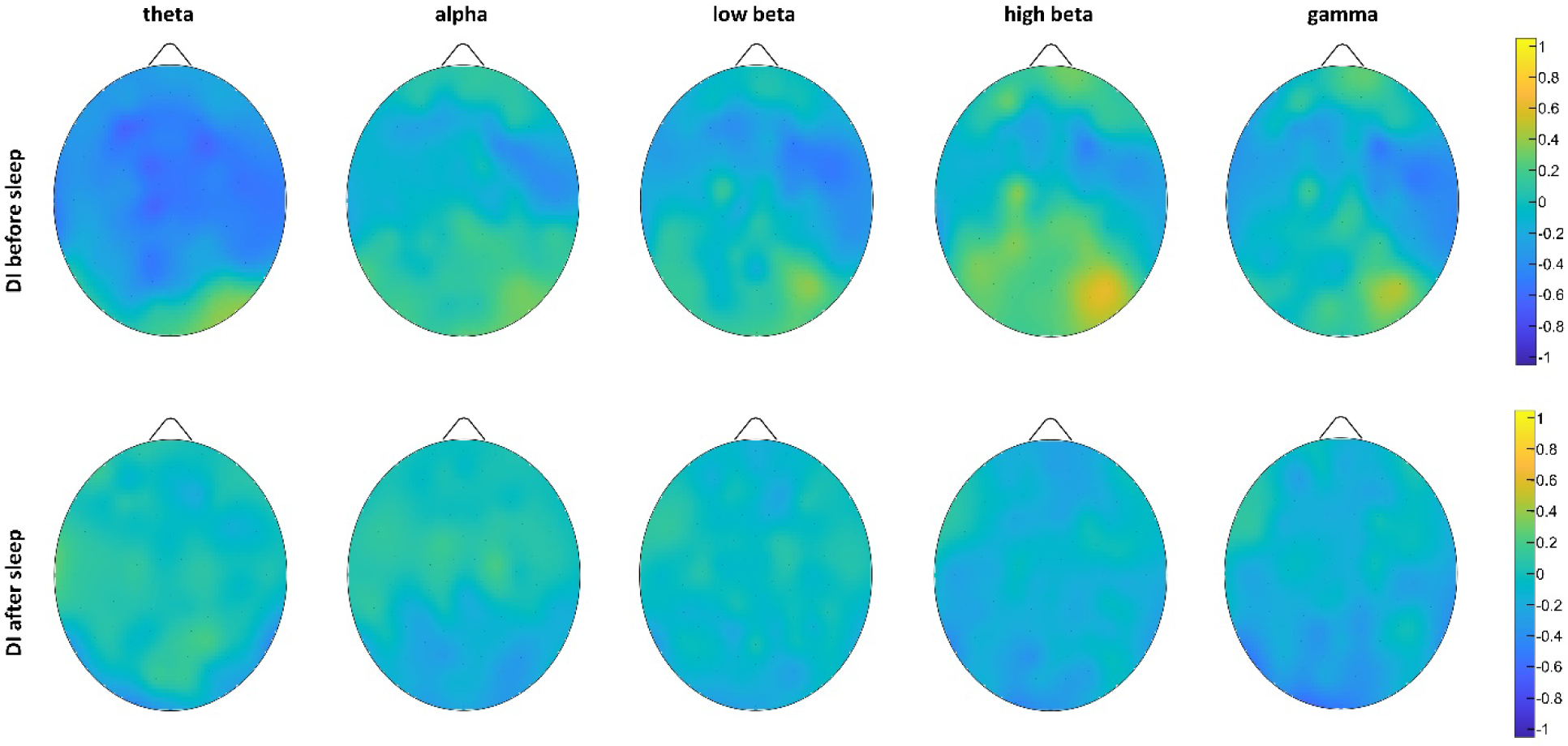
Monocular Deprivation in the morning session (MDmorn): maps of Spearman’s correlations calculated between EEG power band content and deprivation index before and after two hours of dark exposure. No significant correlations between EEG rhythms power and visual plasticity indices were observed.

**Figure S6.**
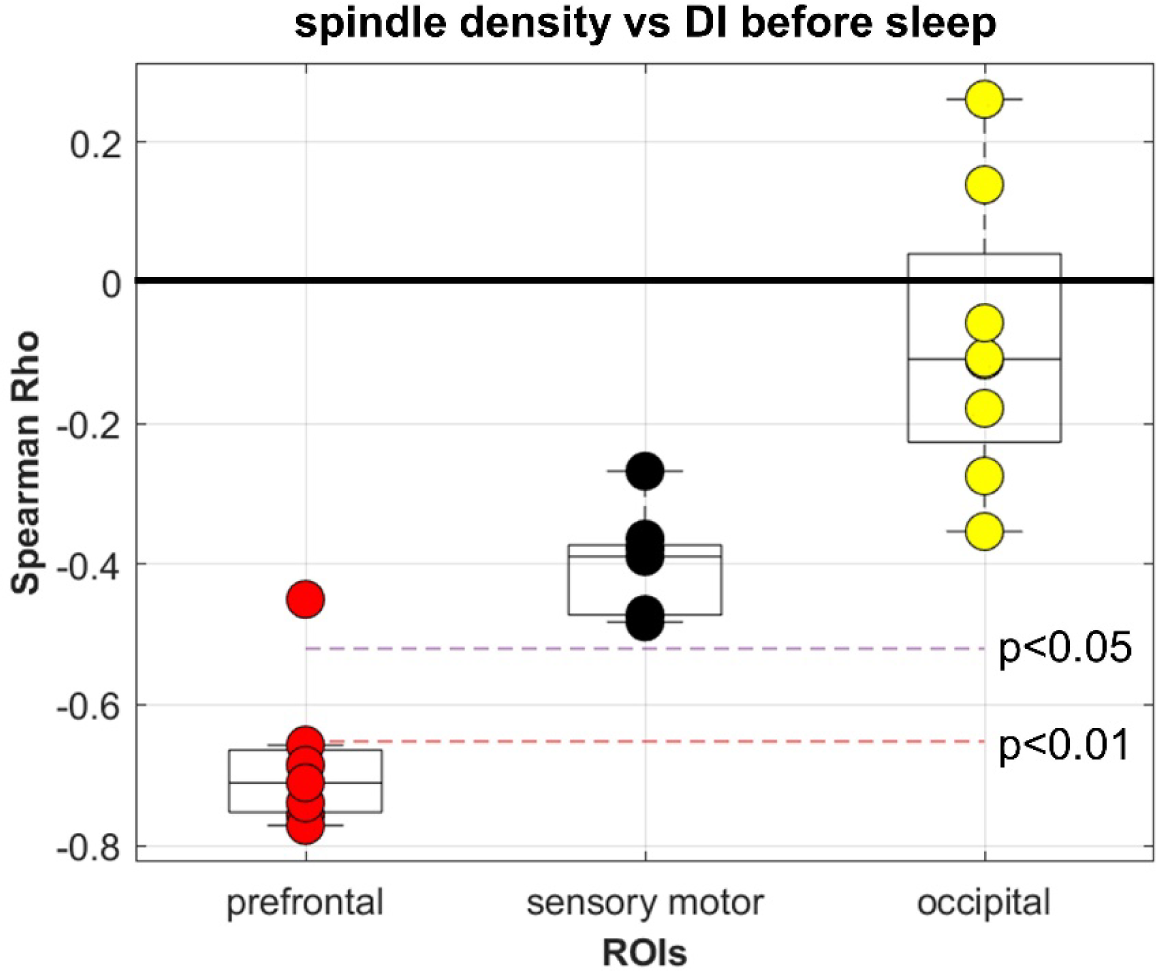
Correlations with DI after sleep. Coherence between electrodes within each ROIs for spindle density (Figure 4). All but one of the electrodes belonging to the prefrontal ROI showed a significant (p>0.01) correlation with DI measured after sleep.

**Figure S7.**
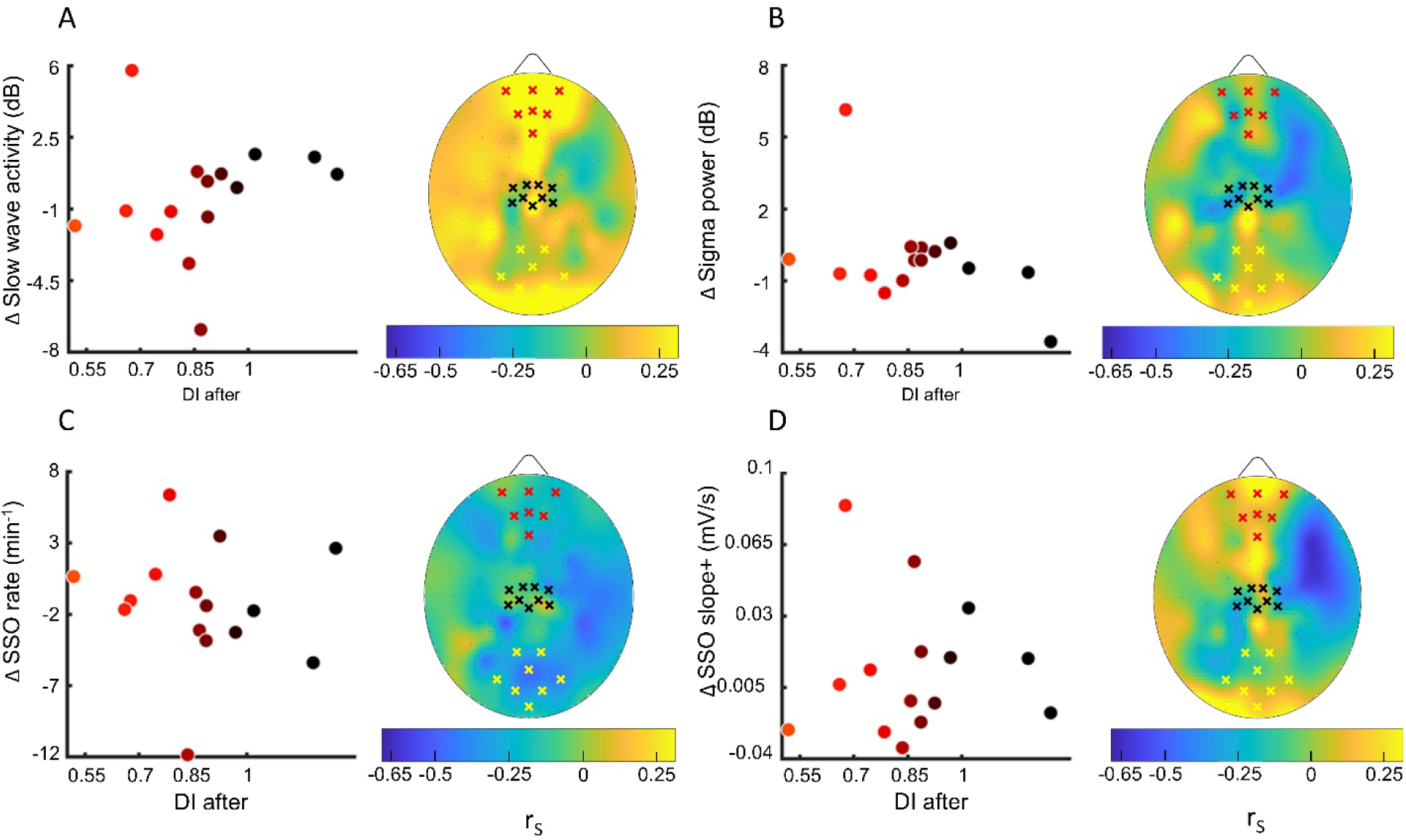
A) A) Correlation between the deprivation index measured **after** sleep (DI after) and the changes from the control night to the MD night of the slow wave activity power averaged over the prefrontal ROI. No electrodes in the defined ROIs (black crosses: sensory-motor control ROI, yellow crosses: occipital ROI, red crosses: prefrontal ROI) shows significant correlation with the deprivation index. Dot’s colour changes from black to red as a function of DI. The scalp map shows the spatial distribution of correlation. . (B) same as A for the sigma activity power; (C) same as A for the occurrence of sleep slow oscillations (SSO); (D) same as A for the steepness of slope+ of SSOs.

**Figure S8.**
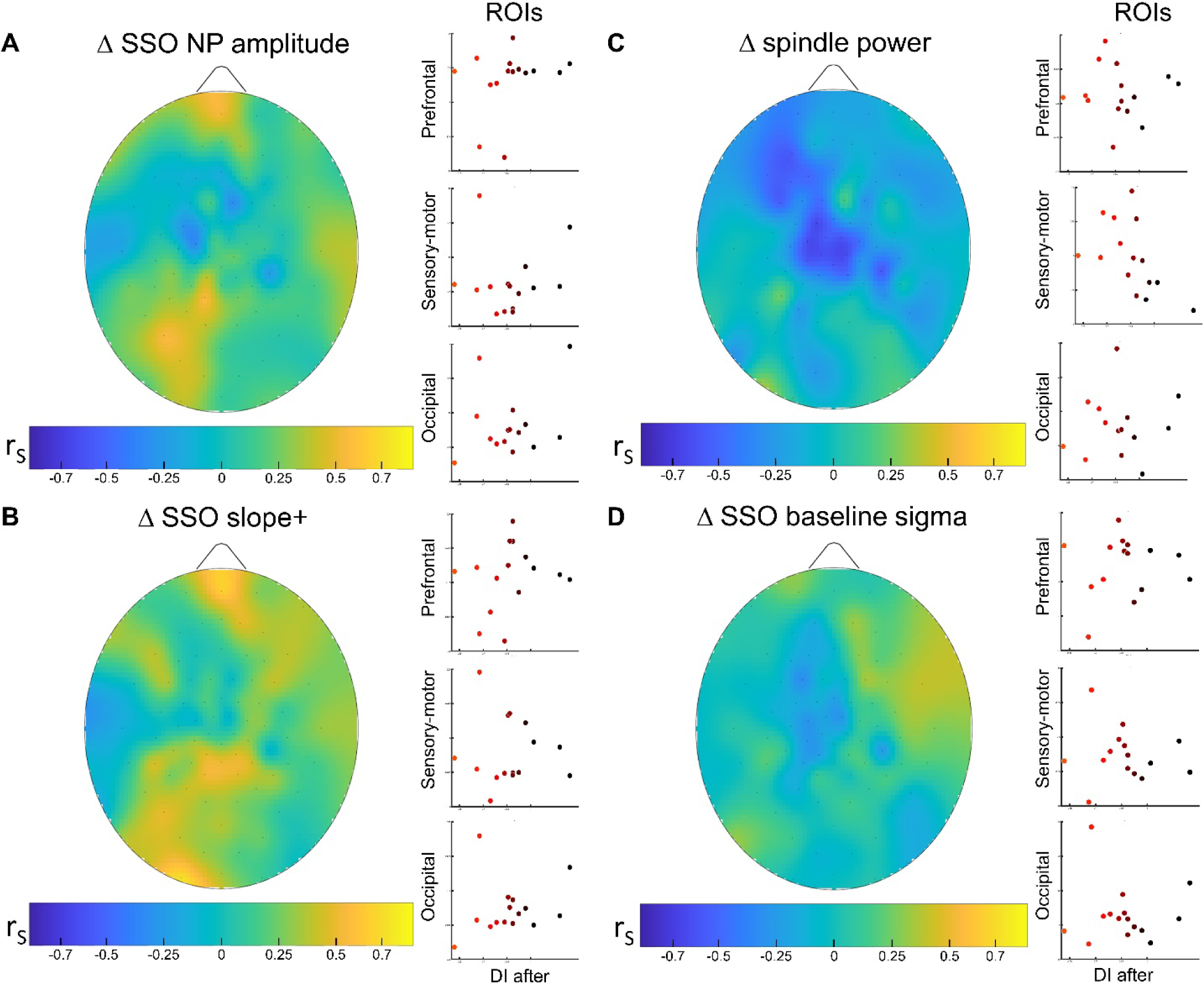
Across sleep analysis. **Correlation analysis between** EEG features changes within the sleep cycle and residual eye dominance plasticity measured at the end of the two hours of sleep. To perform the analysis we 1) partitioned SSO and spindle events into tertiles according to their occurrence, 2) estimated the average measures of events belonging to the first and last tertile, and considered the variation between tertiles as an estimate of the changes across the two hours of sleep. Finally, we tested whether there is a consistent relationship between measures of individual retained plasticity (DI after) and changes in SSO and sleep spindles across the sleep cycle within the three ROIs. None of the parameters considered in the three ROIs have shown significant (after FDR correction) associations with the individual DI after sleep. (A) Correlation between the changes of SSO peak to peak amplitude (NP amp) from the early sleep to the late sleep within the two hours and the DI after. Scatterplot shows individual values averaged within the prefrontal, sensory-motor and occipital ROIs. No significant correlations appeared when considering any ROIs. The scalp map shows the spatial distribution of Spearman’s correlation (rs) for each electrode; (B) same as A for the spindle power; (C) same as A for the steepness of SSO slope+; (D) same as A for the SSO baseline sigma. All other details as Figure S4.

## Supplementary Tables

**Table S1.** Sleep Macrostructural Parameters: MDN-CN descriptive statistics.

|  |  |  |  |  |  |  | Spearman's corr |  |  |  |  |  |
| --- | --- | --- | --- | --- | --- | --- | --- | --- | --- | --- | --- | --- |
|  | MDnight |  | Cnight |  | wilcoxon<br>paired<br>comparison |  | DI before sleep |  |  | DI after sleep |  |  |
|  | median | iqr | median | iqr | p | p_fdr | rho | p | p_fdr | rho | p | p_fdr |
| sleep latency (min) | 7 | 9 | 9 | 10 | 0.62 | 0.86 | 0.14 | 0.62 | 0.70 | 0.15 | 0.59 | 0.91 |
| N2 (min) | 43 | 14 | 43 | 22 | 0.88 | 1.00 | 0.06 | 0.83 | 0.83 | -0.26 | 0.35 | 0.91 |
| N3 (min) | 56 | 23 | 53 | 26 | 0.65 | 0.86 | -0.20 | 0.47 | 0.70 | -0.01 | 0.96 | 0.96 |
| REM (min) | 0 | 8 | 4 | 7 | 0.47 | 0.86 | -0.57 | 0.03 | 0.21 | 0.12 | 0.66 | 0.91 |
| WASO (min) | 6 | 4 | 6 | 5 | 0.28 | 0.86 | -0.22 | 0.43 | 0.70 | 0.12 | 0.68 | 0.91 |
| REM latency (min) | 92 | 44 | 84 | 30 | 0.44 | 0.86 | 0.31 | 0.56 | 0.70 | -0.09 | 0.92 | 0.96 |
| shift phase (1/min) | 0 | 0 | 0 | 0 | 0.28 | 0.86 | -0.23 | 0.42 | 0.70 | -0.41 | 0.13 | 0.50 |
| sleep fragmentation<br>(1/min) | 0 | 0 | 0 | 0 | 1.00 | 1.00 | -0.18 | 0.53 | 0.70 | -0.59 | 0.02 | 0.19 |

iqr= interquartiles range; fdr=FDR correction.

**Table S2:**
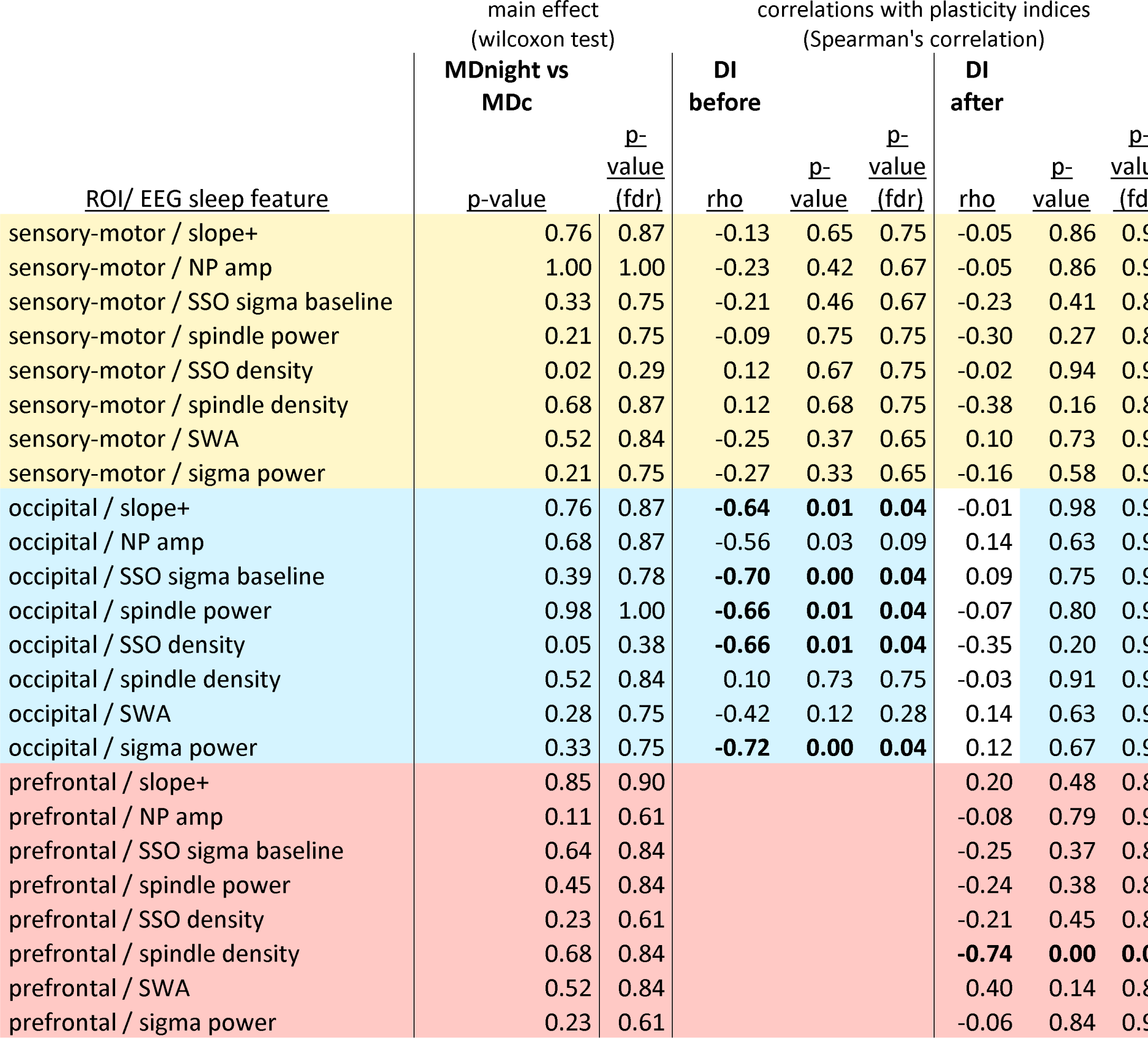
features comparison between conditions and correlations with plasticity indices for the relevant ROIs.

